# Isolated from input: Evidence of default mode network support for perceptually-decoupled and conceptually-guided cognition

**DOI:** 10.1101/150466

**Authors:** Charlotte Murphy, Elizabeth Jefferies, Shirley-Ann Rueschemeyer, Mladen Sormaz, Hao-ting Wang, Daniel S. Margulies, Jonathan Smallwood

## Abstract

The default mode network supports a variety of mental operations such as semantic processing, episodic memory retrieval, mental time travel and mind-wandering, yet the commonalities between these functions remains unclear. One possibility is that this system supports cognition that is independent of the immediate environment; alternatively or additionally, it might support higher-order conceptual representations that draw together multiple features. We tested these accounts using a novel paradigm that separately manipulated the availability of perceptual information to guide decision-making and the representational complexity of this information. Using task based imaging we established regions that respond when cognition combines both stimulus independence with multi-modal information. These included left and right angular gyri and the left middle temporal gyrus. Although these sites were within the default mode network, they showed a stronger response to demanding memory judgements than to an easier perceptual task, contrary to the view that they support automatic aspects of cognition. In a subsequent analysis, we showed that these regions were located at the extreme end of a macroscale gradient, which describes gradual transitions from sensorimotor to transmodal cortex. This shift in the focus of neural activity towards transmodal, default mode, regions might reflect a process of isolation from specific sensory inputs that enables conceptually rich cognitive states to be generated in the absence of input.

**Highlights:**

- Brain regions supporting meaning overlap with stimulus independence.
- Bilateral angular gyri and left MTG respond strongly to both features of cognition.
- These patterns reflect a shift in activity towards regions of transmodal cortex.
- Complex memory representations may emerge in cortical areas distant from input.

## 1. Introduction

Although initial studies characterized the default mode network (DMN) as “task negative”, this network actively supports aspects of cognition (Spreng, 2012), including semantic processing (Binder, Desai, Graves, & Conant, 2009; Krieger-Redwood et al., 2016), episodic recollection (Rugg & Vilberg, 2013), working memory (Konishi, McLaren, Engen, & Smallwood, 2015; Spreng et al., 2014; Vatansever, Menon, Manktelow, Sahakian, & Stamatakis, 2015), autobiographical planning (Spreng, Gerlach, Turner, & Schacter, 2015; Spreng, Stevens, Chamberlain, Gilmore, & Schacter, 2010), self-generation of emotion (Engen, Kanske, & Singer, 2017) and imagining the future or the past (Schacter & Addis, 2007). Although we lack an over-arching account of the functions of the DMN, many of these situations involve memory retrieval – i.e., a requirement to focus cognition on previously-encoded knowledge, as opposed to information in the external environment. In line with this account, many regions within or allied to the DMN are considered to be heteromodal ‘hubs’ for memory-related processes, including the posterior cingulate cortex (Leech, Braga, & Sharp, 2012; Leech & Sharp, 2014), angular gyrus (Bonnici, Richter, Yazar, & Simons, 2016; Seghier, 2013), hippocampus (Moscovitch, Cabeza, Winocur, & Nadel, 2016) and anterior temporal lobes (Visser, Jefferies, & Lambon Ralph, 2010). In addition, cognitive states that activate the DMN tend to involve meaningful content that has personal relevance (Gusnard, Akbudak, Shulman, & Raichle, 2001).

The current study was motivated by the hypothesis that there might be common neurocognitive processes underpinning perceptually-decoupled and conceptually-guided cognition in the DMN. During states of episodic recollection, we recreate past experiences that involve places, objects and people not currently present in the environment. Consequently, memory retrieval might necessitate a process of decoupling from sensorimotor systems, allowing cognition to be generated internally in a way that diverges from what is going on around us (Smallwood, 2013). These perceptually-decoupled states might be easier in brain regions whose neural computations are functionally independent, or distant, from systems important for perceiving and acting. This is consistent with the observation that the distributed regions of the DMN are maximally distant from primary visual and motor cortex, both in terms of their distinct patterns of functional connectivity and their positions along the cortical surface (Margulies et al., 2016).

In addition, DMN regions might support higher-order representations with predictive value across multiple situations and modalities, which integrate features from diverse sensorimotor regions. Contemporary accounts of semantic representation (Lambon Ralph, Jefferies, Patterson, & Rogers, 2017) envisage an interaction between unimodal brain regions that support knowledge about specific features (e.g., knowledge that BANANAS are YELLOW and CURVED in visual cortex) and heteromodal regions within or allied to the DMN, which extract deeper similarity structures across these domains (i.e., allow us to understand that BANANA and KIWI are conceptually related, despite salient differences in colour, shape etc.). This view is also consistent with the observation that DMN lies at the extreme end of a gradient from heteromodal to unimodal cortex (Margulies et al., 2016), since increasingly abstract and complex representations might be formed at greater distances along the gradient, where the influence of specific features linked to stimuli in the immediate environment is reduced (Buckner & Krienen, 2013; Margulies et al., 2016; Mesulam, 1998). Within the DMN, angular gyrus (Binder & Desai, 2011; Bonner et al., 2013) and anterior temporal cortex (Lambon Ralph et al., 2017; Patterson et al., 2007) are both implicated in heteromodal semantic processing. However, their roles remain controversial since other regions such as left inferior frontal gyrus and posterior aspects of the temporal lobe frequently show stronger task-induced activation in fMRI. Angular gyrus, in particular, typically shows a pattern of task-induced deactivation, which is greater for harder judgements in both semantic and non-semantic tasks (Humphreys et al., 2015; Humphreys & Lambon Ralph, 2015). In addition, despite commonalities in the intrinsic connectivity of these regions, differences in semantic content have been proposed although not broadly accepted (Jackson, Hoffman, Pobric & Lambon Ralph, 2016): the anterior temporal lobes might support object identification, while angular gyrus is potentially more sensitive to thematic associations (Davey et al., 2015; Schwartz et al., 2011).

We developed a novel paradigm to identify brain regions important for stimulus independence, more complex memory representations and the combination of both features in cognition. Our experiment builds on prior work by Konishi and colleagues (Konishi et al., 2015). In their study, participants kept track of the location of pairs of simple shapes (triangles, squares and circles) presented either side of fixation. When probed with one shape from a prior trial and asked which side of the screen it was presented on, activity increased in regions including those within the DMN. The current study extended this paradigm by varying the complexity of the information to be encoded and retrieved. In one condition participants keep track of the location of pairs of stimuli that vary on a single feature (colored patches), in a second they keep track of stimuli that vary in a more complex manner (pairs of familiar real world objects such as dogs or cars). Objects place greater demands on memory than do colours because they are distinguished based on a greater number of features. This allowed us to contrast higher and lower levels of representational complexity in the perceptual representations and memories that would be probed. We also manipulated whether these decisions were made when the relevant information is on the screen (0 – back) or when only the identity of the target upon which the decision is based is present (1 – back). In the latter case the relevant spatial information must be retrieved from memory, a manipulation that allowed us to explore the property of stimulus independence in cognition. This paradigm is presented schematically in Figure 1.

**Figure 1.**
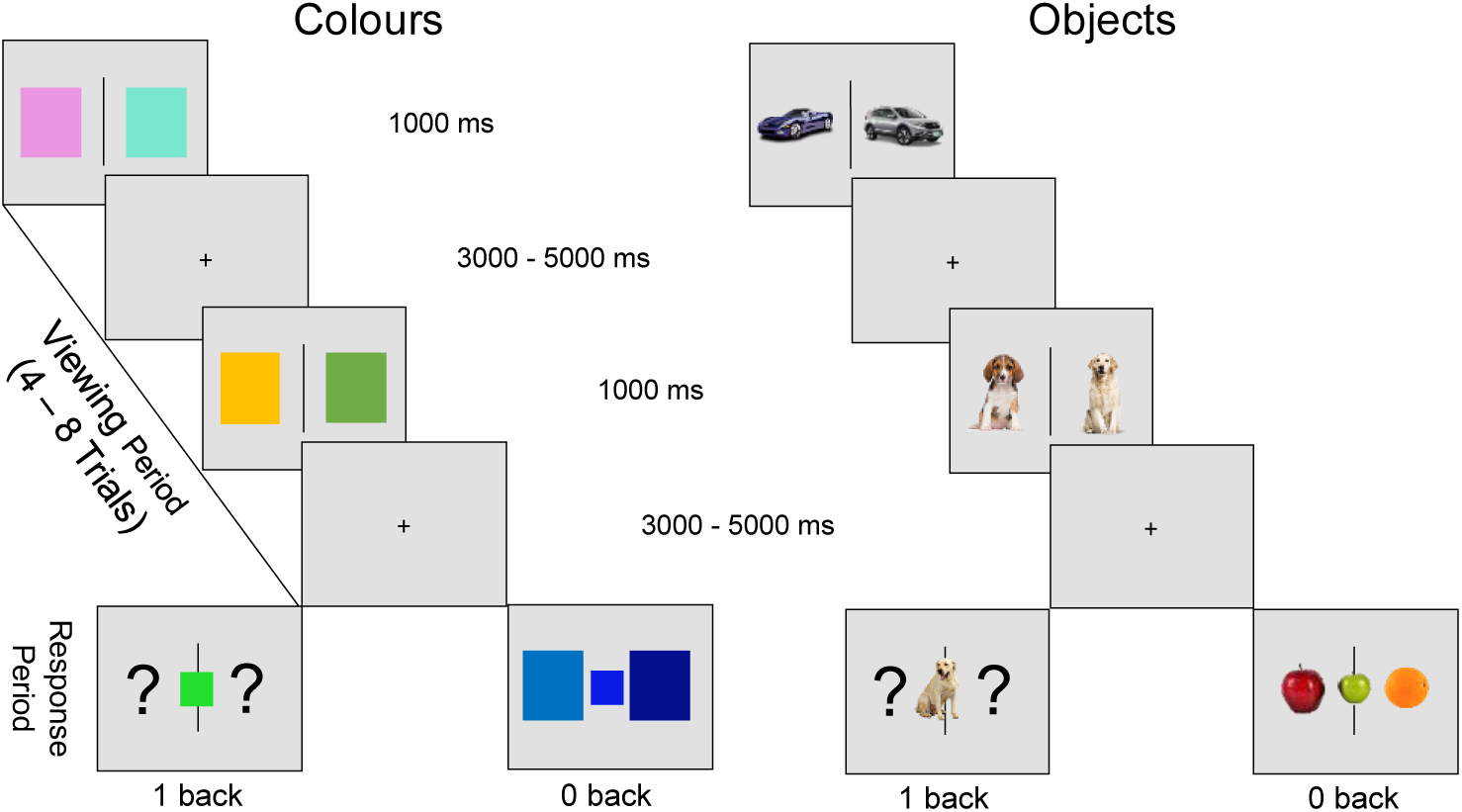
Experimental design. The four different judgments that participants made in this experiment.

We used this paradigm in a functional magnetic resonance imaging (fMRI) experiment to localize the brain regions that support the properties of stimulus independence and representational complexity. Our aim was to establish whether regions sensitive to perceptual decoupling and conceptual retrieval fall within the DMN, and whether these effects were located in overlapping or distinct regions. We identified regions of cortex that respond: (i) to stimuli with a rich multi-modal structure by comparing the response of objects to colors, (ii) when decision making has a higher reliance on memory by comparing decisions that are made in the 1-back condition with those made in the 0-back conditions, and (iii) to conditions that require a combination of both elements of cognition. We tested these hypotheses using both standard whole brain univariate analyses, as well as characterizing neural activity in each condition in terms of its position on the macroscale gradient from unimodal to heteromodal cortex described by Margulies et al. (2016).

If regions in the DMN are activated when spatial decisions are guided by information from memory, this would support a role in stimulus independent decision making. Alternatively, if DMN regions respond to decisions made regarding objects rather than colours, this would reflect a role in the processing of information with a high degree of representational complexity. Finally, if DMN regions show the strongest response when spatial decisions are made based on objects from memory, these regions would support more complex stimulus independent representational states. This latter pattern would be consistent with the hypothesis that decoupling from perceptual input enables cognition to represent information that diverges from what is going on around us (Smallwood, 2013). Moreover, if this processing emerges in regions located towards the transmodal end of the principle gradient, this would support the hypothesis that the “distance” from systems important for perceiving and acting, provides a cortical mechanism that underpins the processing of complex representations derived from memory (Lambon Ralph et al., 2016; Margulies et al., 2016).

## 2. Material and Methods

### 2.1 Participants

Thirty right-handed native British-speaking participants with normal or corrected-to-normal vision were recruited from the University of York (16 female; mean age 22.68, range 18-34 years). One participant’s data was excluded due to excessive motion artefacts, leaving twenty-nine subjects in the final analysis for (15 female; mean age 22.57, range 18-24 years). In a subsequent analysis we used a set of 60 participants’ resting state data, including the same individuals who performed the task (34 female; mean age 20.32, range 18-29 years). Both studies were approved by the York Neuroimaging Centre (YNIC) Ethics Committee. Participant’s provided informed consent prior to the start of the experimental session.

### 2.2 Stimuli

The task paradigm had four conditions: (A) Object 0-back, (B) Object 1-back, (C) Colour 0-back and (D) Colour 1-back using a block design. In all conditions, pairs of items were presented separated by a central line. In the colour conditions, these were different coloured squares, while in the object conditions, these were familiar and meaningful objects, taken from the same semantic category (i.e., different types of cars, fruit, dogs; see Figure 1). Items were presented once with no repetition. The contrast between object and colour conditions allowed us to investigate regions that are important for the retrieval of conceptual information. The colour patches only varied on one feature (their colour), while the objects were meaningful multi-featural concepts. In addition, the contrast of 0-back and 1-back conditions allowed us to investigate the effect of stimulus-independent processing (1 back > 0 back).

### 2.3 Procedure

In the scanner, participants completed a total of four functional runs (average run time 8 min 32 s). Within each run, there were two blocks related to each of the 4 conditions (Object 1-back; Object 0-back; Colour 1-back; Colour 0-back). Each block began with written instructions stating the task type (0-back or 1-back). Blocks consisted of observing pairs of items (1000 ms); each pair was separated by a jittered inter-stimulus interval (ISI; 3000-5000 ms) in which a fixation cross was presented. At random intervals (4-8 trials), a third item was presented in the centre of the screen and participants were asked to indicate the location of one of the pair (left or right) that was most similar to this probe (see Figure 1). This paradigm also required participants to match items that were present and compared this with items in memory. In the 0-back catch-trials participants had to decide which stimulus (left or right of the screen) was most similar to this centrally-presented probe (i.e., all items were present on the screen). In the 1-back catch-trials, participants had to decide which stimulus (left or right of the screen) had been most similar to this centrally-presented probe on the previous trial (i.e., the critical stimulus was absent). Blocks consisted of 5 probes in total and lasted on average 64 s.

### 2.4 MRI Acquisition

Data for both experiments were acquired using a GE 3 T HD Excite MRI scanner at the YNIC. A Magnex head-dedicated gradient insert coil was used in conjunction with a birdcage, radio-frequency insert coil tuned to 127.4 MHz. A gradient-echo EPI sequence was used to collect data from 38 bottom-up axial slices aligned with the temporal lobe (TR = 2s, TE = 18ms, FOV = 192x192mm, matrix size = 64x64, slice thickness = 3mm, slice-gap = 1mm, flip-angle = 90°). Voxel size was 3x3x3mm. Functional images were co-registered onto a T1-weighted anatomical image from each participant (TR = 7.8s, TE = 3ms, FOV = 290x290mm, matrix size = 256x256mm, voxel size = 1.13x1.13x1mm) using linear registration.

### 2.5 Pre-processing

All imaging data were pre-processed using a standard pipeline and analysed via FMRIB Software Library (FSL Version 6.0). Images were skull-stripped using a brain extraction tool [BET, (Smith, 2002)]. The first five volumes (10s) of each scan were removed to minimize the effects of magnetic saturation, and slice-timing correction with Fourier space time-series phase-shifting was applied. Motion correction (MCFLIRT, (Jenkinson, Bannister, Brady, & Smith, 2002)) was followed by temporal high-pass filtering (cut-off = 0.01Hz). Individual participant data was registered to their high-resolution T1-anatomical image, and then into a standard spare (Montreal Neurological Institute); this process included tri-linear interpolation of voxel sizes to 2x2x2 mm.

The resting state functional data used were pre-processed and analysed using the FMRI Expert Analysis Tool (FEAT). The individual subject analysis involved: motion correction using MCFLIRT; slice-timing correction using Fourier space time-series phase-shifting; spatial smoothing using a Gaussian kernel of FWHM 6mm; grand-mean intensity normalisation of the entire 4D dataset by a single multiplicative factor; high-pass temporal filtering (Gaussian-weighted least-squares straight line fitting, with sigma = 100 s); Gaussian low-pass temporal filtering, with sigma = 2.8s

### 2.6 Task based fMRI

For our task-based analysis, the time points of interest were the probe trials where participants had to make a decision about something present (0-back) or absent (1-back) from the screen. We therefore used a box-car regressor to model the probe trial for each condition and another one to model the entire block. Modelling the entire block ensured any effect detected from our analysis can be attributed to the probe itself and not the general effect of the block. Box-car regressors for each probe/block, for each condition, for each run, were convolved with a double gamma hemodynamic response function. Regressors of no interest were included to account for head motion. We computed four contrasts: (1) 0-back > 1-back, (2) 1-back > 0-back, (3) Object > Colour and (4) Colour > Object. A fixed effect design (FLAME, http://www.fmrib.ox.ac.uk/fsl) was conducted to average the four runs, within each individual. Individual participant data were then entered into a higher-level group analysis using a mixed effects design (FLAME, <u>http://www.fmrib.ox.ac.uk/fsl)</u> whole-brain analysis. Finally, our analysis focused on a conjunction of 1-back > 0-back and Object > Colour to identify regions engaged in both stimulus independent processing and conceptually abstract representations.

### 2.7 Resting-state fMRI

We extracted the time series from regions identified by univariate analysis and used these as explanatory variables in a connectivity analyses at the single subject level. In each analysis, we entered 11 nuisance regressors; the top five principal components extracted from white matter (WM) and cerebrospinal fluid (CSF) masks based on the CompCor method (Behzadi, Restom, Liau, & Liu, 2007), six head motion parameters and spatial smoothing (Gaussian) was applied at 6mm (FWHM). WM and CSF masks were generated from each individual’s structural image (Zhang, Brady, & Smith, 2001). No global signal regression was performed, following the method implemented in Murphy, Birn, Handwerker, Jones, & Bandettini (2009).

All whole brain analyses were cluster corrected using a z-statistic threshold of 3.1 to define contiguous clusters. Multiple comparisons were controlled using Gaussian Random Field Theory at a threshold of p < .05 [34]. All statistical maps produced in these analyses are freely available at Neurosynth at the following URL: http://neurovault.org/collections/2296/.

## 3. Results

Table 1 presents behavioural performance, in the form of response efficiency (RT/ACC), for each of the four conditions of our task. Response efficiency controls for speed-accuracy trade-offs. These data were compared using a 2 (task; 0-back vs. 1-back) by 2 (condition; object vs. colour) repeated-measures analysis of variance (ANOVA). There was no significant differences between stimulus type (F(1,28) = 2.55, p = .116) but a significant main effect of task (F(1,28) = 15.38, p < .001). There was no significant interaction (p > .05). These analyses demonstrate that performance on the 1-back task was less efficient than for the 0-back task but that object and colour conditions were well matched in terms of overall task difficulty.

**Table 1.** Behavioural results.

| Condition | Response Efficiency |  |
| --- | --- | --- |
|  | Mean | SE |
| Colour 1-back | 1028 | 206 |
| Colour 0-back | 829 | 287 |
| Object 1-back | 1041 | 229 |
| Object 0-back | 841 | 295 |

Footnote: SE = standard error. Response efficiency = reaction time in milliseconds / percent accuracy.

We next generated statistical maps describing patterns of neural activity at the moments when participants responded in each of our four conditions. These maps were compared at the group level using a GLM (see Methods). The contrast of 0-back > 1-back decisions captures perceptually-guided decision-making, revealing increased activity in the bilateral ventral visual stream, from occipital pole through to posterior fusiform cortex (presented in cool colours in the upper panel of Figure 2). These regions have a well-documented role in online visual processing. The contrast of 1-back > 0-back reflects stimulus independence in decision-making. This comparison revealed greater activation in bilateral angular gyrus and anterior temporal lobes, as well as medial structures in the posterior cingulate cortex and medial prefrontal cortex (these are presented in warm colours in the middle panel of Figure 2). Many of these regions fall within the DMN (58.44% of voxels fell within the DMN as defined by Yeo et al., 2011) and are spatially similar to the ‘general recollection network’ proposed by Rugg and Vilburg (2013). The comparison of Objects > Colours identifies brain areas that support the processing of multi-featural conceptual representations. These are presented in warm colours in the lower panel in Figure 2. This contrast revealed a similar set of regions to the stimulus independence contrast (medial pre frontal cortex, left and right angular gyrus and anterior temporal lobe) with the addition of the right dorsolateral cortex (52.49 % of voxels fell within the DMN as defined by Yeo et al., 2011). The contrast of Colours > Objects yielded no significant whole-brain corrected results. To allow comparison with previous research, the spatial maps for the contrast of 1-back > 0-back from Konishi and colleagues are also displayed: similarities can be seen in posterior cingulate cortex, medial prefrontal cortex, right angular gyrus and dorsolateral cortex.

**Figure 2.**
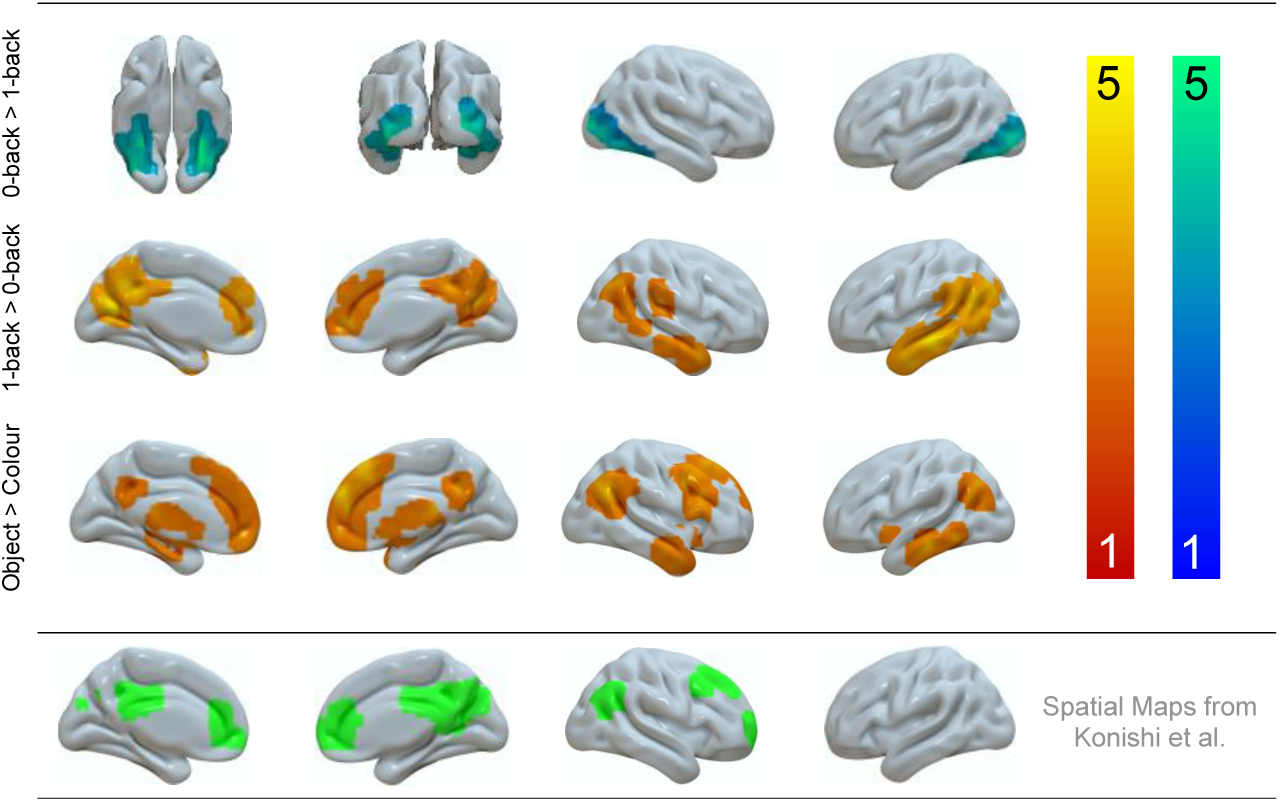
Neural activity produced when making decisions based on meaningful objects and when decisions are made from memory. (a) Activity elicited when decisions were made based using information from perception (b) Activity when decisions were made on the basis of information from memory and (c) when information from memory was more complex. Spatial maps were cluster corrected at Z = 3.1 FWE.

Our next analysis formally identifies regions that show a response to both stimulus independence and memory complexity. Figure 3 shows the results of a formal conjunction of the contrasts of Object > Colour and 1-back > 0-back, revealing three regions – bilateral angular gyrus and lateral medial temporal gyrus in the left hemisphere. The left hand panel of Figure 3 summarizes the parameter estimates from each of these regions in each condition of our task. In every case the strongest response was when decisions were made in the Object 1-back condition. Importantly, although these regions fell within the DMN (88.07% of voxels within the conjunction mask fell within the DMN as defined by Yeo et al., 2011), their response profile indicated greater responding during a demanding condition (i.e. Object 1-back) ruling out a task-negative interpretation of these results.

**Figure 3.**
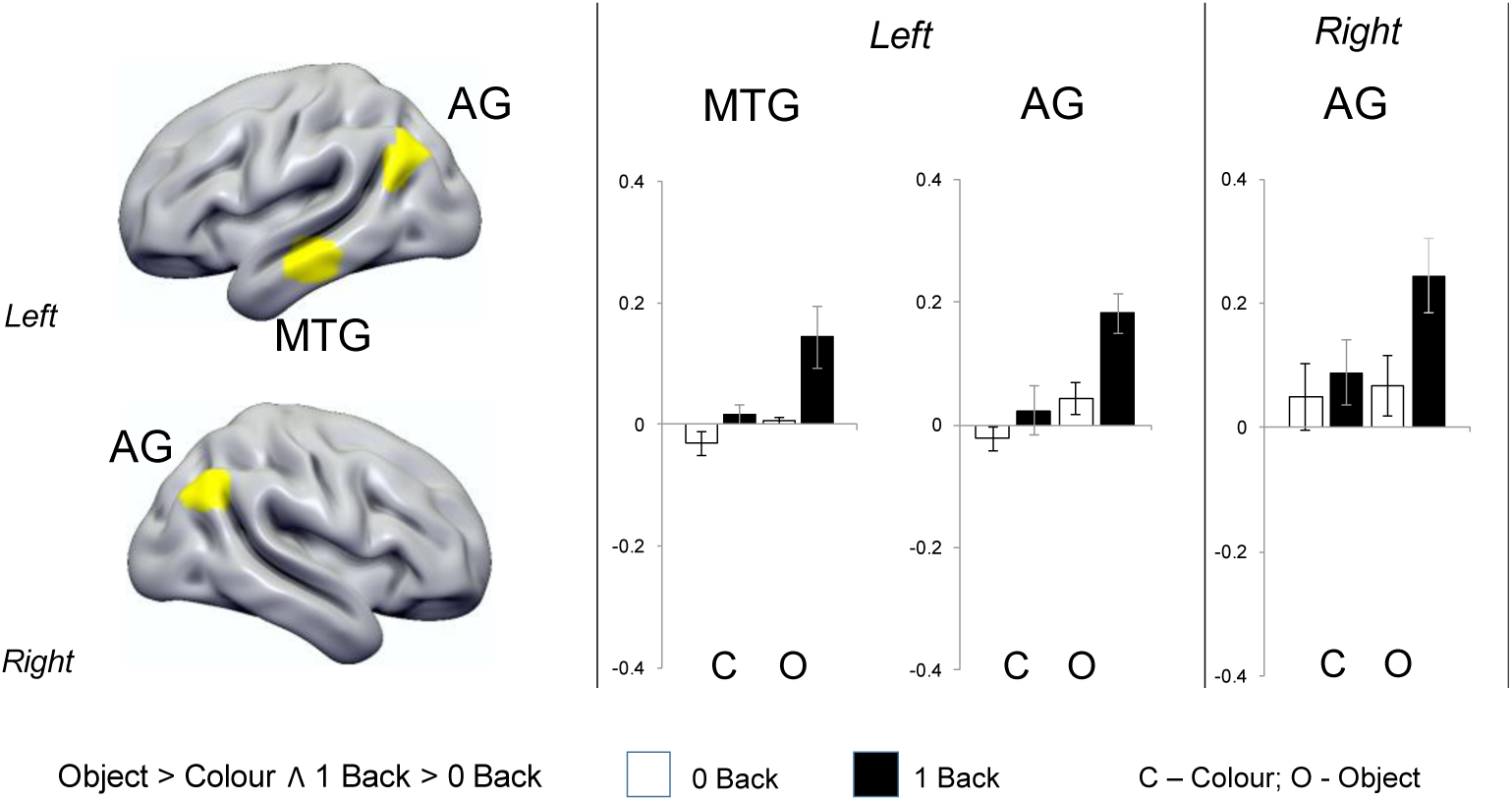
Locating peak activity during stimulus independent decisions regarding complex objects. (a) A conjunction of the neural activity when making decisions based on meaningful categories and when decisions are made in the absence of perceptual input revealed three regions: bilateral angular gyrus and in the left middle temporal gyrus. (b) Percent signal extracted from these regions confirmed an additive effect (i.e. these regions responded significantly more to the object condition when information was not present on the screen compared to all other conditions). The conjunction analysis was based on whole-brain cluster corrected spatial maps from Figure 2. Error bars indicated 95% confidence intervals.

We also explored the intrinsic architecture of conjunction regions responding to Object > Colour and 1-back > 0-back in an independent resting-state data set (see Methods). The results of this analysis are presented in Figure 4 and reveal coupling beyond the seed regions to the posterior cingulate cortex, dorsolateral prefrontal cortex and pre-supplementary motor area bilaterally. Some of these regions fall outside the DMN, as defined by Yeo and colleagues, and instead are members of the frontoparietal network linked with cognitive control (Yeo et al., 2011).

**Figure 4.**
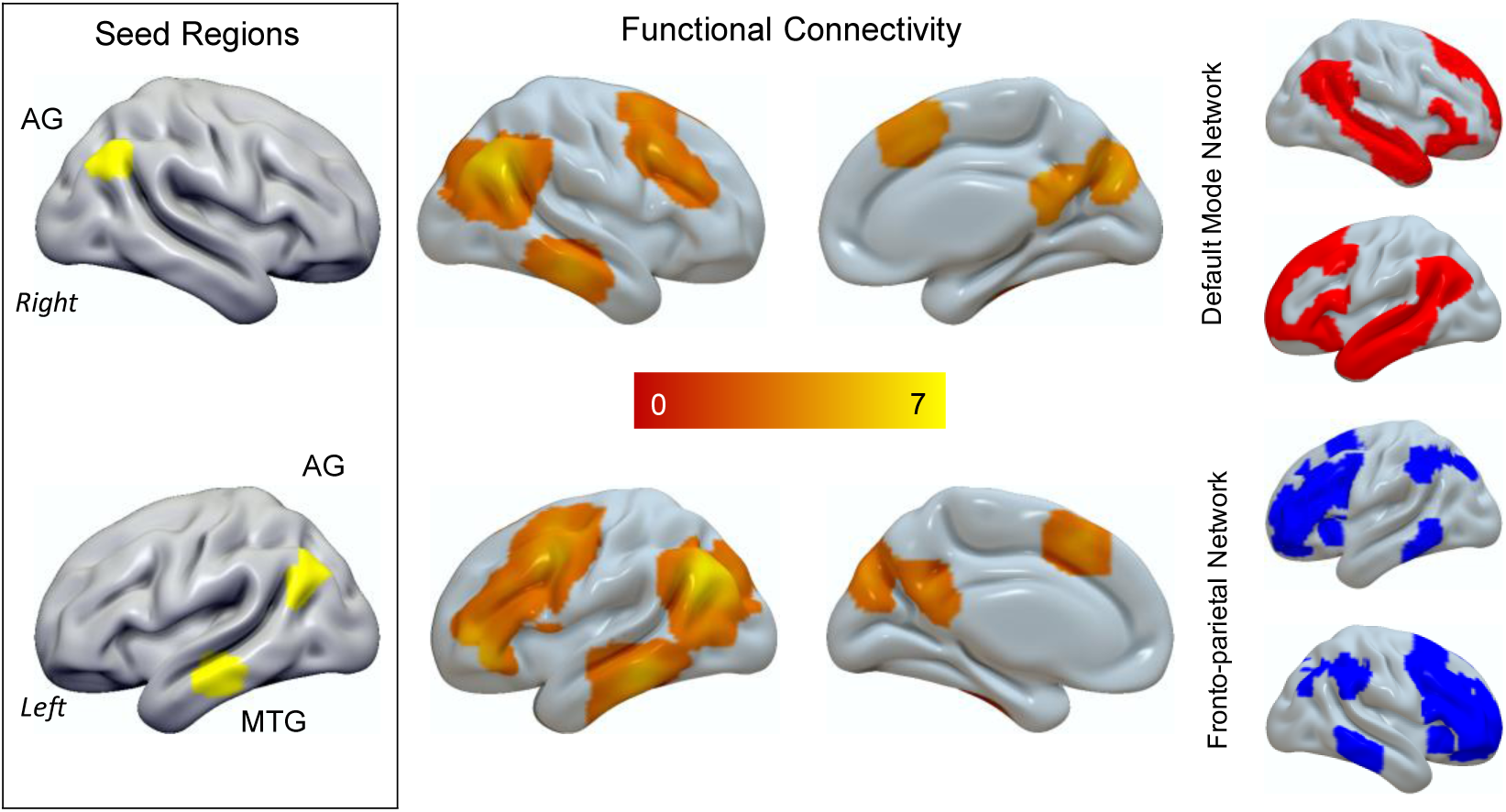
Peak areas during stimulus independent decision regarding complex stimuli involve both regions of the default mode network (DMN) and the fronto-parietal network (FPN). These regions show functional connectivity at rest with both the pre-supplementary cortex and the dorsolateral pre-frontal cortex. Although the regions identified in our conjunction analysis fall within the DMN they show functional communication with regions in the FPN, including the right dorsolateral prefrontal cortex. The spatial networks in the grey panel are from the decomposition of Yeo and colleagues. The conjunction analysis was based on whole-brain cluster corrected spatial maps from Figure 1. For the connectivity analyses spatial maps were cluster corrected at Z = 3.1 FWE.

We also conducted a supplementary analysis contrasting Object and Colour decisions separately in the 1-back and 0-back conditions to confirm regions important for stimulus-independent decisions (see Supplementary Figure 1). This analysis showed that 1-back trials involving meaningful objects activated regions including angular gyrus, middle temporal gyrus and right dorsolateral prefrontal regions more than colours. In contrast, the comparison of Objects > Colours in the 0-back condition only revealed greater activity in fusiform cortex.

Together these analyses highlight a network of regions that are important when spatial decisions are made in the absence of external sensory support, and when they involve multi-feature concepts (Figure 5). Common regions responding to the two task contrasts (1-back > 0-back; Object > Colour), include angular gyrus and middle temporal gyrus. In the right hemisphere, two of the three maps also include the right dorsolateral cortex. All of these right hemisphere regions responded to a similar 1-back > 0-back contrast involving abstract shapes (circle, triangle, square) in the study by Konishi and colleagues (2015).

**Figure 5.**
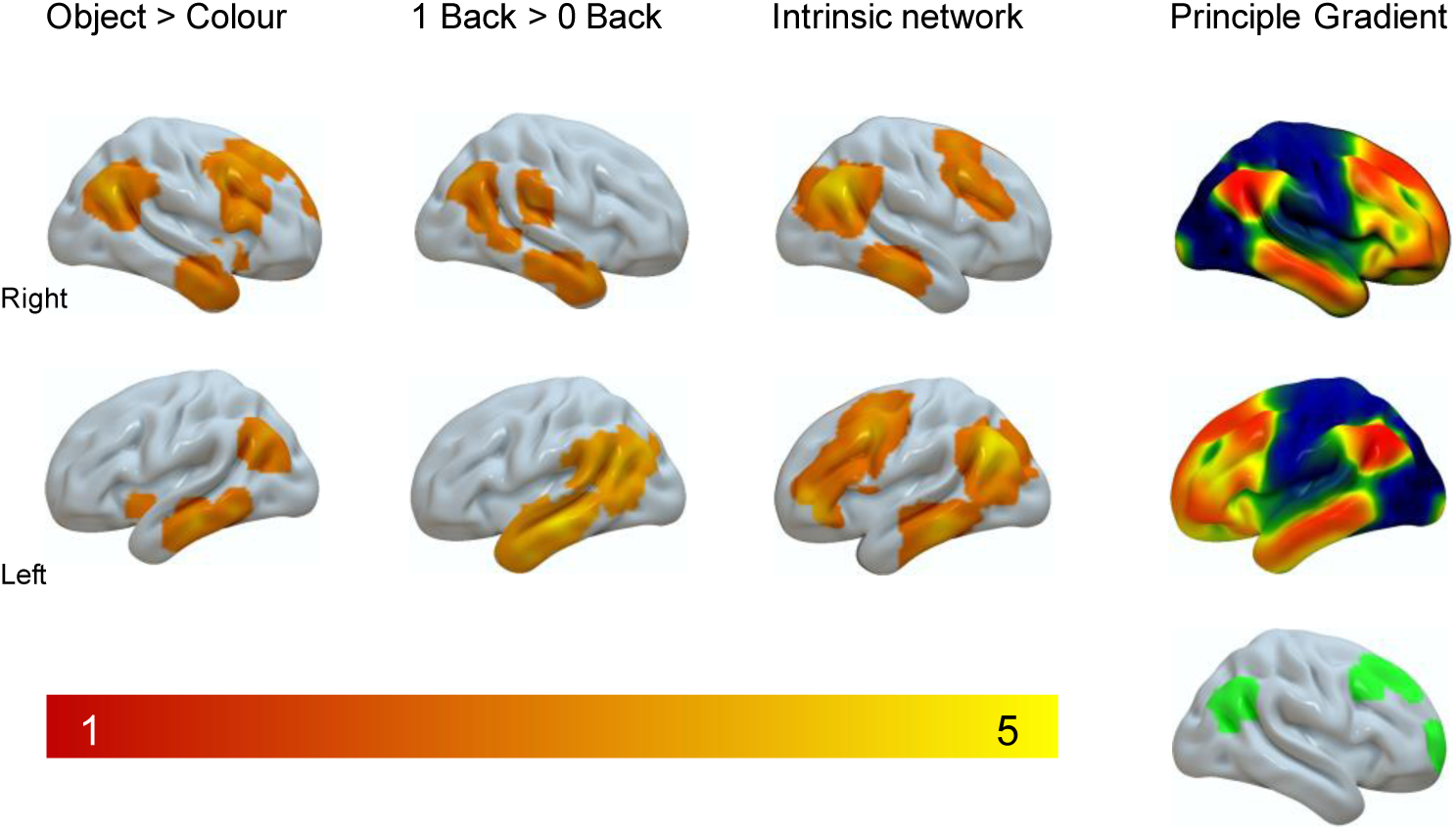
Regions linked to during stimulus independent decisions regarding complex stimuli form localized clusters in transmodal cortex. Regions in the main panel reflect data generated in this experiment. The grey sub panel presents the spatial distribution of the principle gradient from Margulies and colleagues (2016) coloured blue-red, and the cluster corrected map from Konishi and colleagues (2015) coloured green.

In Figure 5, we summarise the spatial maps produced by this experiment and present these alongside the principal gradient from Margulies and colleagues (2016), which describes a functional spectrum of intrinsic connectivity across the cortical surface, extending from primary sensorimotor systems to regions of the DMN at the other extreme. More similar colours on this gradient reflect greater similarity in connectivity. It can be seen that common regions implicated in stimulus-independent and conceptual processing are all localized towards the transmodal end of the principal gradient.

Our final analysis characterizes the similarity between the neural patterns captured by our task and the spatial distribution of the principle gradient from Margulies et al., (2016) in a more formal manner. Following Margulies et al., we divided the principle gradient into 20 equally sized bins. Next for each participant we calculated the average signal in each bin for each condition of our task. The left hand panel in Figure 6 presents these data plotted across the principle gradient separately for each condition; the shaded bars represent the 95% confidence intervals. It can be see that the conditions are most distinct towards the transmodal end, with the highest values when participants made judgments about objects from memory. To quantify these patterns, we compared their distribution using a 2 (stimulus independence) X (stimulus complexity) X 20 (Gradient Bin) ANOVA. This revealed a significant 3-way interaction [F (19, 532) = 5.136, p < .001]. To follow up this interaction, we performed a principle components analysis (PCA) on the condition level data, describing the dynamics captured in the left hand panel of Figure 6. The results revealed two components with eigenvalues greater than 1 accounting for over 86% of the variance (component 1 = 70.49%; component 2 = 15.73%) across the principal gradient bins. The first two components are presented in the right hand panel of Figure 6. The second component describes a gradual transition showing increasing levels of BOLD activity from the unimodal end of the gradient towards the transmodal end. Projecting the values from component 2 back onto the task conditions, and averaging them at the group-level, revealed that this pattern of variance loaded almost exclusively on the ‘object’ 1-back condition. There was a significant positive fit between the spatial map of the principle gradient and recruitment in the Object 1-Back task, but not other conditions.

**Figure 6.**
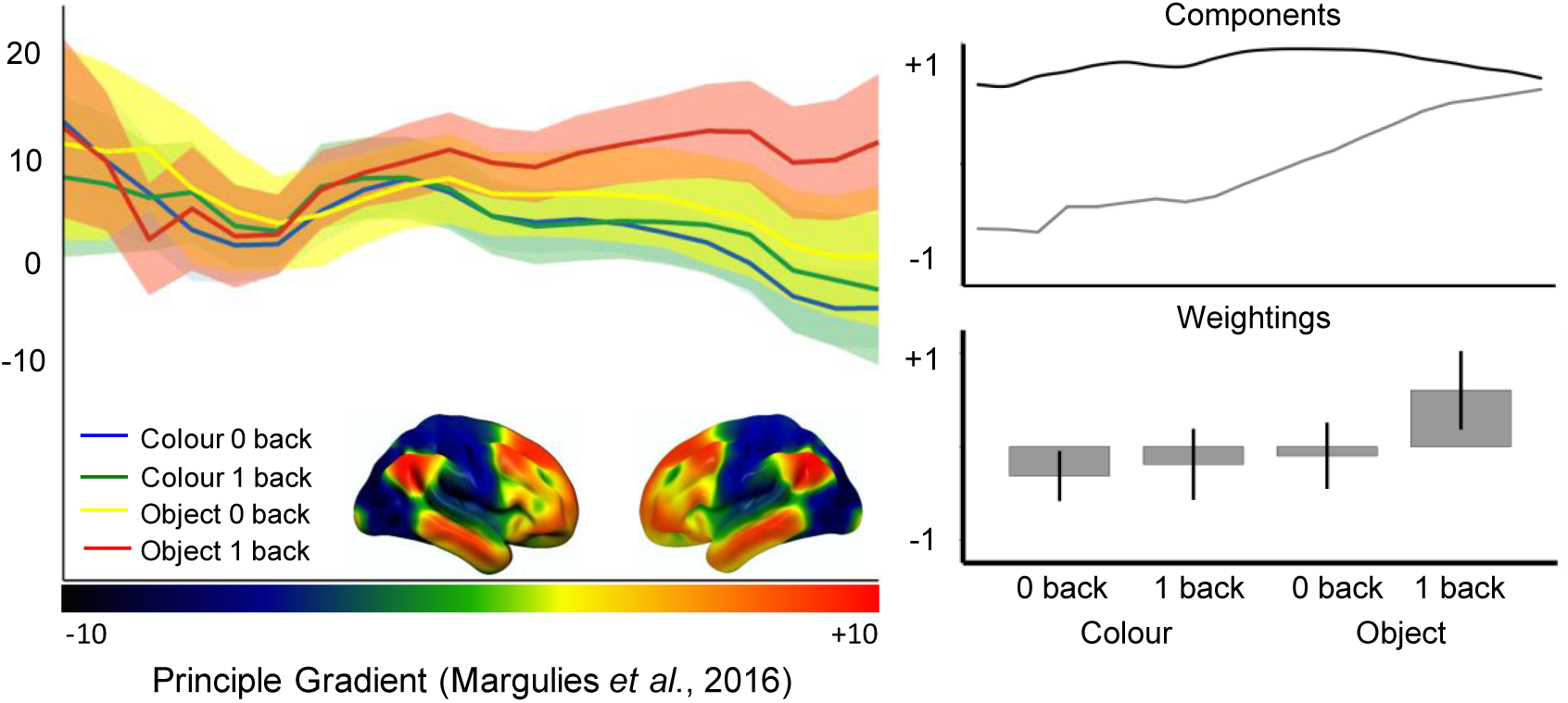
Stimulus independent decisions regarding meaningful objects leads to a whole-brain shift towards the transmodal-end of the gradient. (a) A regions-of-interest analysis using bins of the principal gradient revealed that the decisions that are made on objects rather than colours when these stimuli are not available to perception led to higher activity towards the transmodal-end of the principal gradient. (b) Decomposition using PCA revealed that this difference was related to a gradual shift in the locus of neural activity away from regions on the principal gradient related to perception and action and towards transmodal regions of cortex. Error bars indicated 95% confidence interval.

## 4. Discussion

Our experiment contrasted judgments when stimuli to be decided upon were present in the current trial (stimulus-dependent 0-back decisions) with identical decisions where the information was from the previous trial (stimulus-independent 1-back decisions, see Figure 1). We also varied whether decisions were made on uni-dimensional stimuli (colour patches) or more complex multi-dimensional stimuli (objects in conceptual categories). In each task, the items to be matched were not perceptually identical: participants selected the closest hue or the closest concept from similar distractors. This allowed us to identify regions th**a**t capture cognitive processes important for representing (i) information that is decoupled from stimulus input, (ii) representations that are complex and multi-dimensional in nature and (iii) a combination of both.

Using conventional whole brain analyses we identified overlapping regions in the DMN that are sensitive to both perceptual decoupling (i.e., the requirement to make decisions based on memory, as opposed to the immediate environment) and when these decisions regarded more complex conceptual categories of stimuli (i.e., decisions based on objects rather than colours). We also used a novel analytic approach to demonstrate that these isolated clusters of activity can be seen as part of a whole brain shift in the locus of neural activity towards the extreme end of a gradient from unimodal to heteromodal cortex (Margulies et al., 2016). These findings have broad implications for the role of DMN in cognition, and also contribute to our understanding of specific DMN regions, particularly angular gyrus and lateral temporal lobe. We first consider the results in terms of their implications for functional accounts of these regions. Secondly, we consider the macroscale organisation of the cortex, focusing on approaches which can explain the functional similarity of these distributed clusters and their relative position on the cortical surface.

### Functional implications for the angular gyri

There were stronger responses in left and right angular gyri, as well as in left middle temporal gyrus, when conceptual decisions were based on information that was no longer present in the environment. These findings are inconsistent with several existing accounts of the contribution of angular gyrus to memory and semantic cognition. First, they do not easily align with the proposal that specific aspects of meaning are represented in angular gyrus – namely thematic associations, but not item identity (Davey et al., 2015; Schwartz et al., 2011). Our conceptual task involved matching items on the basis of their identity, yet it robustly activated this region. Secondly, the findings are at odds with the proposal that the angular gyri only activate during contrasts of easier versus harder tasks, and for “automatic” and not “controlled” patterns of retrieval (Humphreys et al., 2015; Humphreys & Lambon Ralph, 2017). The 1-back condition was harder than the 0-back condition and still elicited a greater response.

Instead, our findings are consistent with suggestions that the role of the angular gyri role is to allocate attention to complex memory representations. The angular gyri show a stronger response to a range of memory retrieval situations in which the retrieved representations are detailed, specific or precise (Binder et al., 2005; Price, et al., 2015; Bonnici et al., 2016; Davey at al., 2015). In our study, the 1-back trials required attention to switch from an encoding mode, to a retrieval mode when task relevant information is represented internally. This pattern of responding in the angular gyrus is consistent with the purported role of inferior parietal cortex in focusing attention on memory (Cabeza et al., 2011). In our study, the angular gyri did not activate to the same extent when stimuli were matched on colour, suggesting this region is especially important when heteromodal representations from long-term memory are retrieved.

### Functional implications for temporal cortex

Angular gyrus shows strong intrinsic connectivity with ventral anterior temporal cortex (Davey et al., 2016; Jackson et al., 2016), which is proposed to support the integration of multiple features and modalities to capture ‘deep’ conceptual similarities between items with diverse ‘surface’ features (e.g., items such as PINEAPPLE and KIWI that have different colours, sizes, shapes, phonology etc.; for a review see Lambon (Ralph et al., 2017). Semantic dementia patients with atrophy focussed on this region show highly consistent degradation of conceptual knowledge across tasks (Bozeat et al., 2000; Jefferies & Lambon Ralph, 2006), while neuroimaging studies of healthy participants localise the response during heteromodal conceptual processing to ventral anterior temporal lobes and anterior middle temporal gyrus (Murphy et al., 2017; Visser et al., 2011). Word meaning can be decoded within anterior middle and inferior temporal gyri, while patterns of activation in superior temporal gyrus instead reflect the presentation format (Murphy et al., 2017).

The ventral anterior temporal lobes are thought to provide a “graded hub” in which different unimodal features are gradually integrated to form heteromodal concepts, with visual information reaching this region along the ventral visual pathway (fusiform cortex), auditory and motor information arriving from superior temporal gyrus and frontal cortex, and social/emotional information merging from the temporal pole (Lambon Ralph et al., 2017). Nevertheless, the peak response in the anterior temporal lobes in the current study was in lateral MTG, and not in the site of the putative hub in ventral anterior temporal cortex (Murphy et al., 2017). Visser et al. (2012) observed evidence compatible with two gradients of information convergence in the temporal lobes: first, there is a posterior-to-anterior axis: posterior temporal lobe regions proximal to visual and auditory cortex show largely unimodal responses, while more anterior regions integrate across these types of input to support heteromodal conceptual processing. Secondly, there may be integration from superior and inferior regions, implicated in auditory and visual processing respectively: towards middle temporal gyrus response become more heteromodal response along the length of the temporal cortex. The site we observed in the conjunction of semantic and perceptually-decoupled decisions in the current study corresponds to the extreme heteromodal end of *both* of these temporal lobe gradients.

### Implications for the default mode network

We replicated prior demonstrations that transmodal regions in the DMN are engaged when participants make decisions that rely on information from memory rather than input from perception, even though the 1-back task was more difficult than the 0-back task (Konishi et al., 2015). This pattern of task-positive behaviour adds to a growing body of evidence that the DMN contributes in an active manner to demanding external cognitive tasks (Konishi et al., 2015; Krieger-Redwood et al., 2016; Spreng et al., 2014; Spreng et al., 2015; Spreng et al., 2010; Vatansever et al., 2015). The contribution of DMN to controlled cognitive states appears to reflect situations in which DMN regions work in tandem with the frontoparietal network. Prior work has established the combination of these networks is important for tasks including controlled semantic retrieval (Krieger-Redwood et al., 2016), working memory (Vatansever et al., 2015), autobiographical planning (Spreng et al., 2014; Spreng et al., 2015), retrieving memories of close personal friends (de Caso et al., 2017) and the control of spontaneous thoughts in a deliberate manner (Golchert et al., 2017). Our study shows that right angular gyrus, within the DMN, and right dorsolateral prefrontal cortex, a member of the frontoparietal network, activate together when participants make judgments about meaningful objects from memory rather than colours (see Supplementary Figure 1). Our functional connectivity analysis also demonstrates that these regions are correlated at rest, suggesting they form an intrinsic network. The right dorsolateral cluster replicates the spatial distribution observed from the prior study by Konishi et al. (Konishi et al., 2015) and overlaps with a region of greater grey matter associated with more deliberate mind-wandering (Golchert et al., 2017). Both 1-back retrieval in our paradigm, and more deliberate spontaneous thought, require memory retrieval to be shaped in a goal-directed fashion. It is possible that a range of states requiring the goal-directed control of memory depend on co-operation between these two large-scale networks.

At the most general level our study supports the hypothesis that the capacity for complex memory representations to influence cognition emerges from the topographical arrangement of neural processes across the cortex. Prior work highlighted regions of transmodal cortex, such as the default mode network, as having the greatest distance from uni-modal sensorimotor cortex in both functional and structural space (Margulies et al., 2016). Our finding builds on this observation by showing an association between ongoing neural activity and this dimension of connectivity under situations when the demands placed on cognition require a combination of memory complexity and stimulus independency. Using both standard and novel methods of analysis, we demonstrated that the neural activity associated with this type of activity is prevalent in transmodal regions (Figure 5) and can be represented as a whole brain shift in the balance of neural activity, away from sensorimotor regions cortex and towards the transmodal end of the gradient (Figure 6). This topographical shift in the distribution of neural processing is consistent with theoretical accounts that assume that more abstract cortical functions are facilitated through functional isolation from incoming input (Buckner & Krienen, 2013; Margulies et al., 2016; Mesulam, 1998; Smallwood, 2013). Consistent with this interpretation of DMN function, a recent study found that strong connectivity within the DMN (including an overlapping region of left temporal cortex) was linked to poor performance on tasks which depend on encoding information from the environment but not for those that depended on retrieving information from memory (Poerio et al., 2017). The findings of Poerio and colleagues, in combination with those from the current study, provide converging evidence that regions of the DMN support a state where cognition is guided by memory rather than input, regardless of whether it is beneficial to the task or not.

There are a number of limitations that should be borne in mind when considering the results of this study. First, our comparison of semantic and colour decisions allowed us to demonstrate a neural pattern associating conceptual processing with stimulus independency. This comparison is too crude a manipulation to determine which aspects of the semantic judgements gave rise to this response in the DMN; for example, is it the richness of concepts such as Labrador or apple, their heteromodal nature or the fact that they are acquired over a lifetime, which dissociates them from colours? Future studies could probe different features of retrieval, such as whether the target is a concrete or abstract concept, whether it has to be identified at a specific or superordinate level, and whether there are differences according to the modality of the representation being probed. Second, the nature of our design precludes the ability to separate different aspects of memory retrieval engaged during 1-back decisions. In our paradigm, these decisions require both the integration of appropriate information from memory, as well as the inhibition of the memory representation for the non-probed item. Interestingly, studies have implicated dorsolateral prefrontal cortex in the suppression of memories (Anderson et al., 2004) whereas the angular gyrus has been linked to the integration of appropriate semantic features (Wagner et al., 2015) and the retrieval of specific information (Davey et al., 2015). It is possible that the angular gyrus and dorsolateral prefrontal region are performing distinct roles in integration of relevant associations and suppression of irrelevant information during retrieval in our paradigm. Future work could address this question by manipulating the level of featural overlap between target and probe during retrieval in a similar paradigm as in this experiment.

### Funding

The research was supported by BBSRC grant BB/J006963/1. EJ was supported by a grant from the European Research Council (SEMBIND - 283530). JS was supported by a grant from the European Research Council (Wandering Minds – 303701).

---

**Figure S1.**
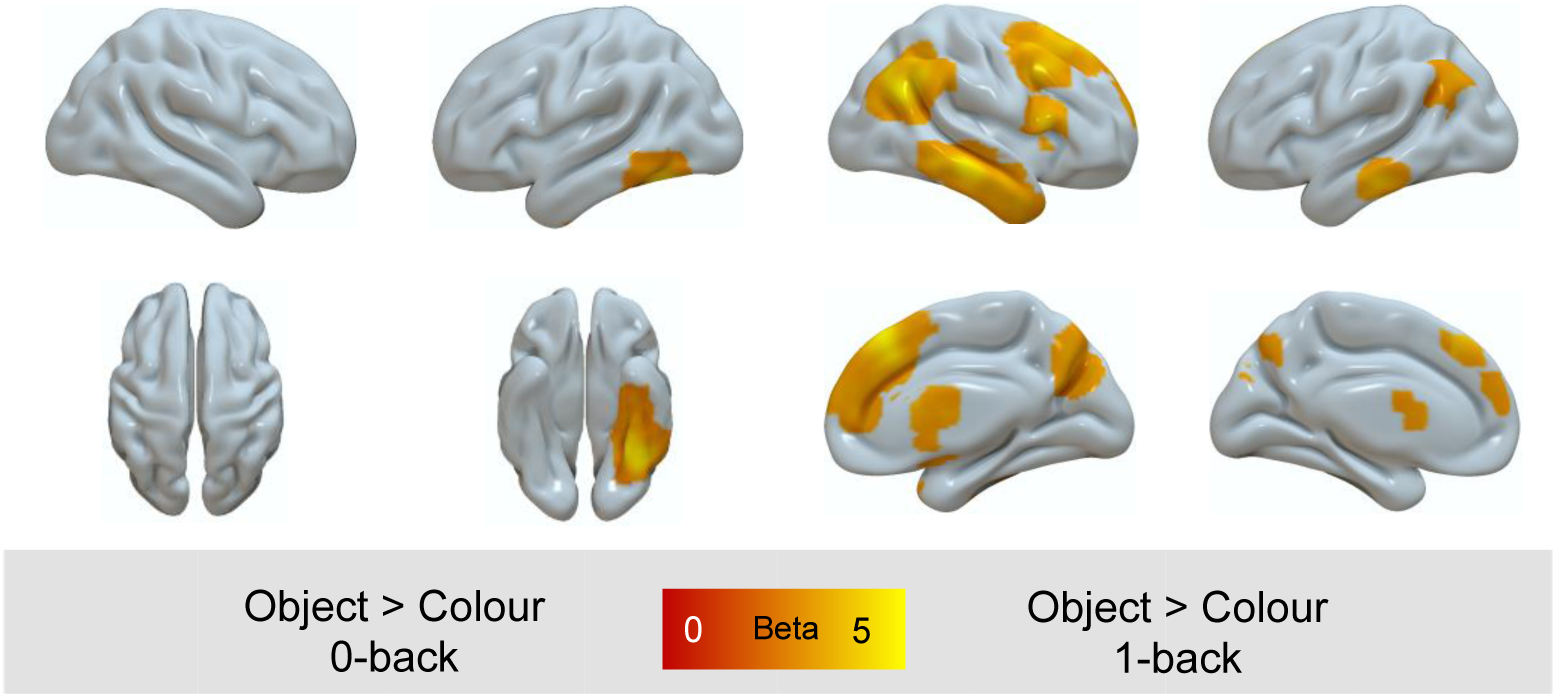
Comparison of complex memory representation in the presence or absence of relevant perceptual input. Spatial maps were cluster corrected at Z = 3.1 FWE.

## References

Anderson, M. C., Ochsner, K. N., Kuhl, B., Cooper, J., Robertson, E., Gabrieli, S. W., … Gabrieli, J. D. (2004). Neural systems underlying the suppression of unwanted memories. Science, 303(5655), 232–235. doi:10.1126/science.1089504

Behzadi, Y., Restom, K., Liau, J., & Liu, T. T. (2007). A component based noise correction method (CompCor) for BOLD and perfusion based fMRI. Neuroimage, 37(1), 90–101. doi:10.1016/j.neuroimage.2007.04.042

Binder, J. R., Desai, R. H., Graves, W. W., & Conant, L. L. (2009). Where is the semantic system? A critical review and meta-analysis of 120 functional neuroimaging studies. Cereb Cortex, 19(12), 2767–2796. doi:10.1093/cercor/bhp055

Bonnici, H. M., Richter, F. R., Yazar, Y., & Simons, J. S. (2016). Multimodal Feature Integration in the Angular Gyrus during Episodic and Semantic Retrieval. J Neurosci, 36(20), 5462–5471. doi:10.1523/JNEUROSCI.4310-15.2016

Buckner, R. L., & Krienen, F. M. (2013). The evolution of distributed association networks in the human brain. Trends Cogn Sci, 17(12), 648–665. doi:10.1016/j.tics.2013.09.017

Cabeza, R., Mazuz, Y. S., Stokes, J., Kragel, J. E., Woldorff, M. G., Ciaramelli, E., … Moscovitch, M. (2011). Overlapping parietal activity in memory and perception: evidence for the attention to memory model. J Cogn Neurosci, 23(11), 3209–3217. doi:10.1162/jocn_a_00065

Davey, J., Cornelissen, P. L., Thompson, H. E., Sonkusare, S., Hallam, G., Smallwood, J., & Jefferies, E. (2015). Automatic and Controlled Semantic Retrieval: TMS Reveals Distinct Contributions of Posterior Middle Temporal Gyrus and Angular Gyrus. J Neurosci, 35(46), 15230–15239. doi:10.1523/JNEUROSCI.4705-14.2015

de Caso, I., Karapanagiotidis, T., Aggius-Vella, E., Konishi, M., Margulies, D. S., Jefferies, E., & Smallwood, J. (2017). Knowing me, knowing you: Resting-state functional connectivity of ventromedial prefrontal cortex dissociates memory related to self from a familiar other. Brain Cogn, 113, 65–75. doi:10.1016/j.bandc.2017.01.004

Engen, H. G., Kanske, P., & Singer, T. (2017). The neural component-process architecture of endogenously generated emotion. Soc Cogn Affect Neurosci, 12(2), 197–211. doi:10.1093/scan/nsw108

Golchert, J., Smallwood, J., Jefferies, E., Seli, P., Huntenburg, J. M., Liem, F., … Margulies, D. S. (2017). Individual variation in intentionality in the mind-wandering state is reflected in the integration of the default-mode, fronto-parietal, and limbic networks. Neuroimage, 146, 226–235. doi:10.1016/j.neuroimage.2016.11.025

Gusnard, D. A., Akbudak, E., Shulman, G. L., & Raichle, M. E. (2001). Medial prefrontal cortex and self-referential mental activity: relation to a default mode of brain function. Proc Natl Acad Sci U S A, 98(7), 4259–4264. doi:10.1073/pnas.071043098

Jenkinson, M., Bannister, P., Brady, M., & Smith, S. (2002). Improved optimization for the robust and accurate linear registration and motion correction of brain images. Neuroimage, 17(2), 825–841.

Konishi, M., McLaren, D. G., Engen, H., & Smallwood, J. (2015). Shaped by the Past: The Default Mode Network Supports Cognition that Is Independent of Immediate Perceptual Input. PLoS One, 10(6), e0132209. doi:10.1371/journal.pone.0132209

Krieger-Redwood, K., Jefferies, E., Karapanagiotidis, T., Seymour, R., Nunes, A., Ang, J. W., … Smallwood, J. (2016). Down but not out in posterior cingulate cortex: Deactivation yet functional coupling with prefrontal cortex during demanding semantic cognition. Neuroimage, 141, 366–377. doi:10.1016/j.neuroimage.2016.07.060

Leech, R., Braga, R., & Sharp, D. J. (2012). Echoes of the brain within the posterior cingulate cortex. J Neurosci, 32(1), 215–222. doi:10.1523/JNEUROSCI.3689-11.2012

Leech, R., & Sharp, D. J. (2014). The role of the posterior cingulate cortex in cognition and disease. Brain, 137(Pt 1), 12–32. doi:10.1093/brain/awt162

Margulies, D. S., Ghosh, S. S., Goulas, A., Falkiewicz, M., Huntenburg, J. M., Langs, G.,… Smallwood, J. (2016). Situating the default-mode network along a principal gradient of macroscale cortical organization. Proc Natl Acad Sci U S A, 113(44), 12574–12579. doi:10.1073/pnas.1608282113

Mesulam, M. M. (1998). From sensation to cognition. Brain, 121 (Pt 6), 1013–1052.

Moscovitch, M., Cabeza, R., Winocur, G., & Nadel, L. (2016). Episodic Memory and Beyond: The Hippocampus and Neocortex in Transformation. Annu Rev Psychol, 67, 105–134. doi:10.1146/annurev-psych-113011-143733

Ralph, M. A., Jefferies, E., Patterson, K., & Rogers, T. T. (2017). The neural and computational bases of semantic cognition. Nat Rev Neurosci, 18(1), 42–55. doi:10.1038/nrn.2016.150

Rugg, M. D., & Vilberg, K. L. (2013). Brain networks underlying episodic memory retrieval. Curr Opin Neurobiol, 23(2), 255–260. doi:10.1016/j.conb.2012.11.005

Schacter, D. L., & Addis, D. R. (2007). The cognitive neuroscience of constructive memory: remembering the past and imagining the future. Philos Trans R Soc Lond B Biol Sci, 362(1481), 773–786. doi:10.1098/rstb.2007.2087

Seghier, M. L. (2013). The angular gyrus: multiple functions and multiple subdivisions. Neuroscientist, 19(1), 43–61. doi:10.1177/1073858412440596

Smallwood, J. (2013). Distinguishing how from why the mind wanders: a process-occurrence framework for self-generated mental activity. Psychol Bull, 139(3), 519–535. doi:10.1037/a0030010

Smith, S. M. (2002). Fast robust automated brain extraction. Hum Brain Mapp, 17(3), 143–155. doi:10.1002/hbm.10062

Spreng, R. N. (2012). The fallacy of a “task-negative” network. Front Psychol, 3, 145. doi:10.3389/fpsyg.2012.00145

Spreng, R. N., DuPre, E., Selarka, D., Garcia, J., Gojkovic, S., Mildner, J., … Turner, G. R. (2014). Goal-congruent default network activity facilitates cognitive control. J Neurosci, 34(42), 14108–14114. doi:10.1523/JNEUROSCI.2815-14.2014

Spreng, R. N., Gerlach, K. D., Turner, G. R., & Schacter, D. L. (2015). Autobiographical Planning and the Brain: Activation and Its Modulation by Qualitative Features. J Cogn Neurosci, 27(11), 2147–2157. doi:10.1162/jocn_a_00846

Spreng, R. N., Stevens, W. D., Chamberlain, J. P., Gilmore, A. W., & Schacter, D. L. (2010). Default network activity, coupled with the frontoparietal control network, supports goal-directed cognition. Neuroimage, 53(1), 303–317. doi:10.1016/j.neuroimage.2010.06.016

Vatansever, D., Menon, D. K., Manktelow, A. E., Sahakian, B. J., & Stamatakis, E. A. (2015). Default mode network connectivity during task execution. Neuroimage, 122, 96–104. doi:10.1016/j.neuroimage.2015.07.053

Visser, M., Jefferies, E., & Lambon Ralph, M. A. (2010). Semantic processing in the anterior temporal lobes: a meta-analysis of the functional neuroimaging literature. J Cogn Neurosci, 22(6), 1083–1094. doi:10.1162/jocn.2009.21309

Wagner, I. C., van Buuren, M., Kroes, M. C., Gutteling, T. P., van der Linden, M., Morris, R. G., & Fernandez, G. (2015). Schematic memory components converge within angular gyrus during retrieval. Elife, 4, e09668. doi:10.7554/eLife.09668

Yeo, B. T., Krienen, F. M., Sepulcre, J., Sabuncu, M. R., Lashkari, D., Hollinshead, M., … Buckner, R. L. (2011). The organization of the human cerebral cortex estimated by intrinsic functional connectivity. J Neurophysiol, 106(3), 1125–1165. doi:10.1152/jn.00338.2011

Zhang, Y., Brady, M., & Smith, S. (2001). Segmentation of Brain MR Images Through a Hidden Markov Random Field Model and the Expectation-Maximization Algorithm. IEEE Transactions on Medical Imaging, 20(1), 45–57.

